# Chromoplast differentiation in bell pepper (*Capsicum annuum*) fruits

**DOI:** 10.1101/2020.09.29.299313

**Authors:** Anja Rödiger, Birgit Agne, Dirk Dobritzsch, Stefan Helm, Fränze Müller, Nina Pötzsch, Sacha Baginsky

**Affiliations:** Plant Biochemistry, Institute of Biochemistry and Biotechnology, Martin-Luther-Universität Halle-Wittenberg, Halle (Saale), Germany; Biochemistry of Plants, Biology and Biotechnology, Ruhr-University Bochum, Bochum, Germany; Biochemistry and Functional Proteomics, Institute of Biology II, University of Freiburg, Freiburg, Germany

**Keywords:** quantitative proteomics, CN PAGE, plastid differentiation, chromorespiration

## Abstract

We report here a detailed analysis of the proteome adjustments that accompany chromoplast differentiation from chloroplasts during bell-pepper fruit ripening. While the two photosystems are disassembled and their constituents degraded, the cytochrome b_6_f complex, the ATPase complex as well as Calvin cycle enzymes are maintained at high levels up to fully mature chromoplasts. This is also true for ferredoxin (Fd) and Fd-dependent NADP reductase, suggesting that ferredoxin retains a central role in the chromoplasts redox metabolism. There is a significant increase in the amount of enzymes of the typical metabolism of heterotrophic plastids such as the oxidative pentose phosphate pathway (OPPP), amino acid and fatty acid biosynthesis. Enzymes of chlorophyll catabolism and carotenoid biosynthesis increase in abundance, supporting the pigment reorganization that goes together with chromoplast differentiation. The majority of plastid encoded proteins declines but constituents of the plastid ribosome and AccD increase in abundance. Furthermore, the amount of plastid terminal oxidase (PTOX) remains unchanged despite a significant increase in phytoene desaturase (PDS) levels, suggesting that the electrons from phytoene desaturation may be consumed by another oxidase. This may be a particularity of non-climacteric fruits such as bell pepper, that lack a respiratory burst at the onset of fruit ripening.

## Introduction

Plastids are semiautonomous organelles that develop and differentiate from undifferentiated proplastids into different plastid types that are distinguished by their pigment content and energy metabolism, for example as photosynthetic, autotrophic or non-photosynthetic, heterotrophic. Because of their relevance for fruit ripening and endosperm development, amyloplasts and chromoplasts received most consideration among the non-photosynthetic plastid types lately (Camara et al., 1995, Waters and Pyke 2005, Sadali et al., 2019). Chromoplasts synthesize large amounts of presumably health-promoting carotenoids, therefore, research on chromoplast differentiation is dedicated to advance carotenoid biosynthesis in crops at a quantitative scale to improve their dietary quality (Taylor and Ramsay 2005, Rodriguez-Concepcion et al., 2018). Chromoplast differentiation can be reversible such as in pumpkin and citrus fruits or irreversible such as in bell pepper and tomato. Irreversible chromoplast differentiation in the latter two species originates from fully functional chloroplasts and proceeds to mature chromoplasts by degradation of thylakoid membranes and their replacement with storage structures for carotenoids (Camara et al., 1995, Egea et al., 2010).

The transition from photosynthetic chloroplasts to fully differentiated heterotrophic chromoplasts requires the remodeling of an autotrophic metabolism to a heterotrophic metabolism and as such represents an integral part of the metabolic reorganizations that occur during fruit ripening (Li and Yuan 2013, Pesaresi et al. 2014). Details of this reorganization at the protein/enzyme level were initially mapped by quantitative proteome analyses in tomato, a climacteric fruit that undergoes a steep rise in respiration and ethylene production at the onset of ripening (Barsan et al., 2012, Szymanski et al., 2017, Quinet et al., 2019). Using three stages during chromoplast differentiation, a decrease in the amount of thylakoid membrane components and carbohydrate metabolic enzymes and a rise in the abundance of stress-response proteins and carotenoid biosynthetic enzymes was reported for the chloroplast to chromoplast transition (Barsan et al., 2010, Barsan et al., 2012, Suzuki et al., 2015). Overall, main portions of the chromoplast proteome match the occurrence of major metabolic modules in other heterotrophic plastid types such as etioplasts and amyloplasts (e.g. Neuhaus and Emes, 2000, Kleffmann et al., 2007, Renato et al., 2015, Ma et al., 2018).

A distinctive feature of chromoplasts compared to other heterotrophic plastid types is the massive synthesis and storage of carotenoids. Recently a DnaJ-like chaperone termed ORANGE (OR) was reported to promote carotenoid biosynthesis by increasing the amount of carotenoid biosynthetic enzymes through functional interactions with the stromal Clp protease complex (Rodriguez-Concepcion et al., 2019). OR promotes the correct folding of phytoene synthase (PSY) and such increases its activity and prevents misfolding and removal by the Clp protease complex, which is similar to the anticipated function of ClpB3 in the stabilization of deoxyxylulose 5-phosphate synthase (DXS) (D’Andrea and Rodriguez-Concepcion, 2019). Furthermore, OR promotes the formation of carotenoid-sequestering structures, and also prevents carotenoid degradation. Intriguingly, OR is also nuclear-localized where it interacts with the transcription factor TCP14 to repress chloroplast biogenesis during de-etiolation suggesting that OR may in general be relevant for plastid-type transitions (Sun et al., 2019). A posttranscriptional mechanism that could be important for chromoplast differentiation was recently suggested to comprise the remodeling of the protein import machinery by the E3 ligase Sp1 (Ling et al., 2012). In this model, the ubiquitin proteasome system (UPS) disassembles the Toc159-containing TOC complexes that sustain the import of photosynthetic proteins in chloroplasts to TOC complexes with different specificities to promote the import of chromoplast-specific proteins (Sadali et al., 2019).

We report here a comprehensive proteome analysis of the chromoplast differentiation process in the non-climacteric fruit bell pepper. We performed native PAGE to separate protein complexes from seven different stages during chromoplast differentiation and performed *in solution* analyses with the same fractions. Using protein quantification by MSE-based Hi3 method (Silva et al., 2006, Helm et al., 2014), we quantified proteins in every fraction in three biological replicates. Our study provides a detailed quantitative map of chromoplast differentiation at the protein level and is available as a high resolution resource for plant biologists.

## Results and Discussion

### Isolation of plastids from bell pepper fruits

*Selma Bell,* the bell pepper variety used in this study, produces full-sized green fruits within approximately two weeks after flowering, representing the initial stage of fruit ripening. About one week later the fruit color changes into dark-green, almost black. Three to four weeks later, the fruits reach the second intermediate stage with mixed red and green colors. Within one week after this point, the fruits turn homogenously red and are fully matured, representing the final stage of fruit development (Figure 1A and 1B). These stages were used for the isolation of plastids from ripening fruit tissue by an isolation procedure that is based on Percoll density gradient centrifugation (Siddique et al., 2006). The employed step-gradient allowed the separation of plastid populations that differ by their density, distributing into an upper and a lower band (Fig. 1). All plastid-containing bands were collected, named by their occurrence in the gradient as e.g. GU for green-upper band, IL for intermediate-lower band and so on and their pigment content was determined by HPLC combined with UV/VIS spectra measurements that were acquired in the range of 260-760 nm. Using a monitoring wavelength of 430 nm, 15 peaks could be numbered in order of their occurrence, as shown in Supplemental Figure S1. LC-APCI-MS was used to determine m/z values of these peaks. Due to the combination of UV/VIS and m/z, 11 peaks could be identified (Supplemental Fig. S1, Supplemental Table S1).

**Figure 1:**
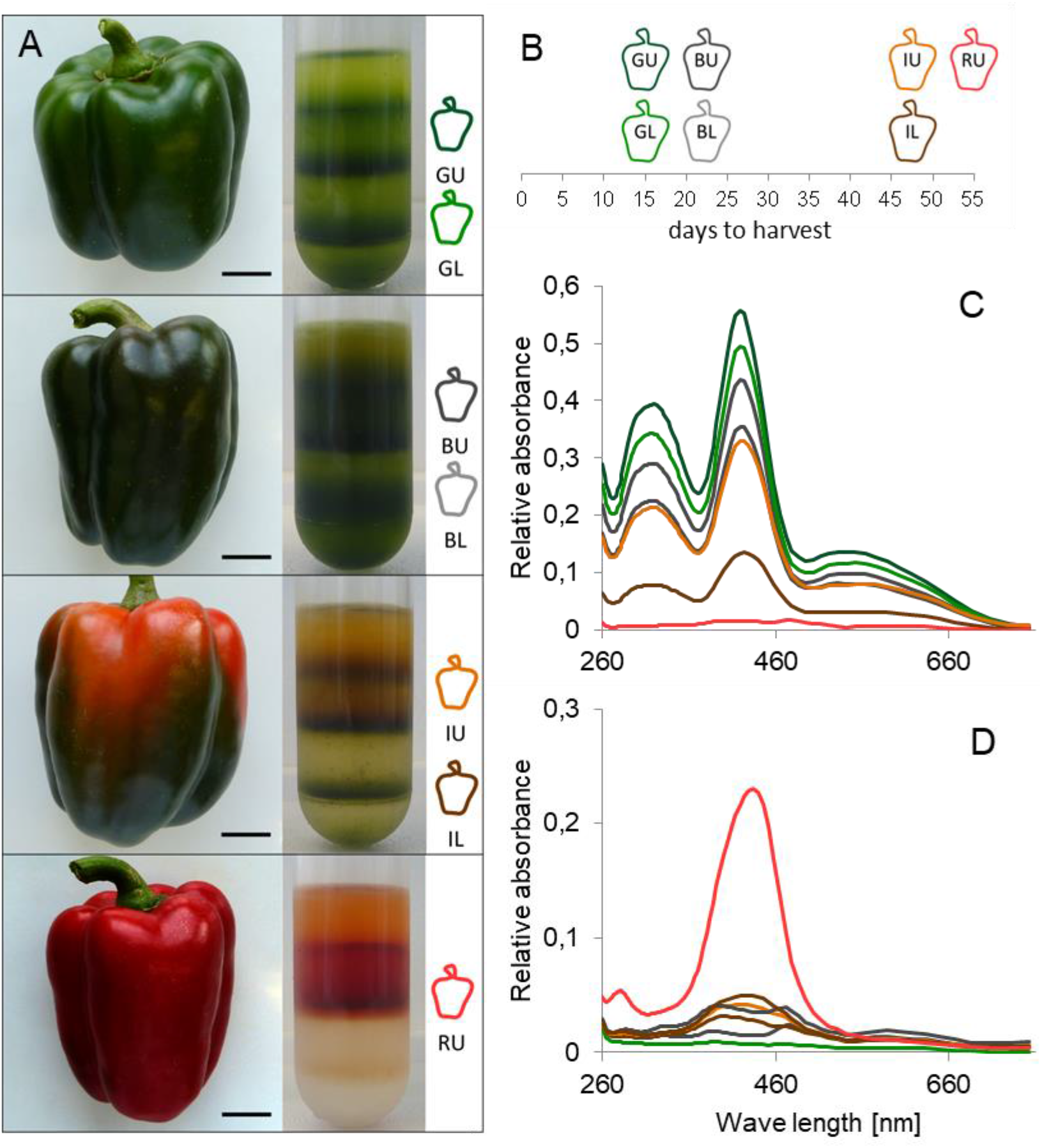
Bell pepper fruit ripening. (A) Bell pepper fruits harvested at different time points during fruit ripening are presented along with the corresponding Percoll gradients obtained after density gradient centrifugation of their homogenates. (B) Time-scale in days (0 representing flowering time) for the harvest, illustrated by pepper icons for the different developmental stages. All stages of maturation are represented: GL, GU, BL, BU, IL, IU, RU. (B + C) Kinetic of changes in pigment content are illustrated for the chloroplast specific chlorophyll b peak (C), and the chromoplast specific peak of the carotenoid diester capsanthin-dimyristate (D). Abbreviations: G – green, B – black, I – intermediate, R – red, L – lower. U – upper.

The differentiation of chromoplasts from chloroplasts is accompanied by the degradation of chlorophyll and the synthesis of carotenoids. We therefore assessed the pigment content of the different plastid preparations by exploring chlorophyll b (Figure 1C, Peak 5 in Supplemental Fig. S1), and capsanthin-dimyristate (Figure 1D, peak 7 Supplemental Fig. S1) abundance. A progressive decline in chlorophyll b levels among the different preparations supported the intermediate nature of the black and red/green fruits. Together with the increase in carotenoid levels, the pigment measurements provided the following order to the different plastid preparations on their way to chromoplasts: starting from fully developed chloroplasts in GU/GL, BU, BL, IU, IL, to RU as the final stage of chromoplast differentiation. The kinetics of the capsanthin-dimyristate peak shows that the most fundamental changes in carotenoid content occur relatively late, i.e. between IU/IL and RU. This suggests that carotenoid biosynthesis and sequestration have their maximum during the final days of fruit ripening. Consistently, we identified several compounds characteristic for fully ripe red fruits, e.g. capsanthin/capsorubin-laurate, zeaxanthin dipalmitate as well as zeaxanthin laurate myristate exclusively at the last maturation stage (Supplemental Fig. S1, Supplemental Table S1, peak 11-13). In contrast, there was a slow decline in chlorophyll content suggesting that chlorophyll degradation occurs throughout the entire chromoplast differentiation process.

### Protein complex reorganization during progressive chromoplast differentiation

We extracted proteins from the seven different plastid preparations and subjected the protein fractions to colorless native gel electrophoresis (CN-PAGE). The distribution of protein complexes supports progressive chromoplast differentiation in the order inferred by the pigment measurements (see above, Fig. 1 and Fig. 2). To determine the identity of the visualized protein complexes, we identified and quantified proteins from the GL and the RU lanes from 16 gel pieces by MSE analyses (Silva et al., 2006, Helm et al., 2014) (Fig. 2, Supplemental Fig. S2). The abundant complexes visible in the GL lane comprise PSII and PSI subunits that decrease in abundance from GL/GU to BU, IL and IU. In the RU stage, these complexes are no longer visible (Fig. 2). This preparation is dominated by prominent low-molecular mass protein bands that appear approximately at stage IL.

**Figure 2:**
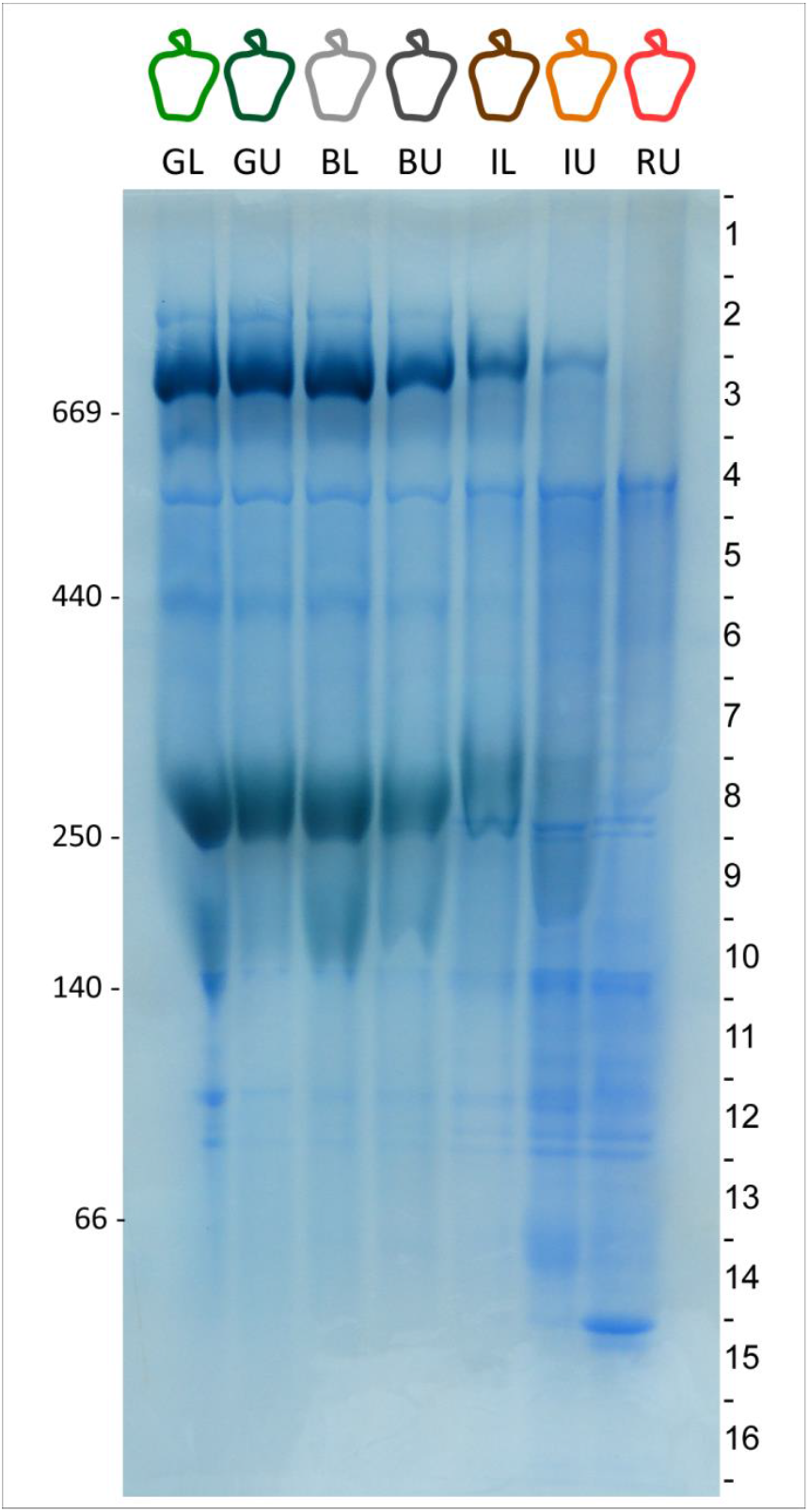
Native protein complexes during chromoplast differentiation. Colorless Native PAGE of extracts from isolated plastids obtained from the different developmental stages as detailed in Fig. 1B. The gel was stained with Coomassie and subsequently cut into 16 pieces for tryptic digest and mass spectrometric protein identification, as indicated.

For the purpose of a rapid automated functional annotation by MapMan (Thimm et al., 2004), we matched the identifications to the *Arabidopsis thaliana* database (TAIR10) and provide the identity of individual proteins in the different gel fractions in Supplemental Table S2. Initial screening of the quantitative data suggested, that the cytochrome b_6_f complex, ATPase and other electron carriers (bin 1.1.5) are sustained from chloroplasts to fully developed chromoplasts (Supplemental Fig. S2). Enzymes involved in carotenoid biosynthesis are significantly upregulated, together with plastoglobuli-associated proteins for carotenoid sequestration. Reproducibility of protein quantification in proteomics approaches suffers from large numbers of experimental interventions. We therefore decided to move ahead with our analysis by performing a single-step *in solution* digest with the seven plastid preparations detailed above, and inferring proteome adaptations during chromoplast differentiation from these data (see below).

### Proteome abundance dynamics during progressive chromoplast differentiation

Protein extracts from the plastid samples obtained at different ripening stages were compared by their protein patterns on SDS-PAGE (Figure 3A). While the chloroplast samples are dominated by low molecular mass light-harvesting complex proteins, the chromoplast samples accumulate increasing amount of capsanthin/capsorubin synthase (CCS) at approximately 50 kDa (Siddique et al., 2006). This enzyme is the most abundant protein in chromoplasts supporting the highly active carotenoid biosynthesis in this plastid type (AT3G10230, Supplemental Tables S2 and S3). The decrease of the LHC band at 25-30 kDa and the increase of the 50 kDa CCS band supports a progressive transition between the plastid types represented by our samples (Fig. 3 A). We used these fractions for an *in solution* digest and performed quantitative proteomics by matching the MS-data against the Uniprot bell pepper proteome database (UP000222542) using an internal abundance marker for Hi3-based absolute protein quantification (see also above, Silva et al., 2006, Helm et al., 2014). Altogether we performed three biological replicates with three technical replicates each, giving rise to a cumulative identification of 2537 proteins after removal of spurious detections (<1 fmol sum in all fractions), histone proteins and cytosolic ribosome contaminations. Surprisingly, only very few mitochondrial contaminations were detected that were confined to the upper band in every preparation (Supplemental Table S3).

**Figure 3:**
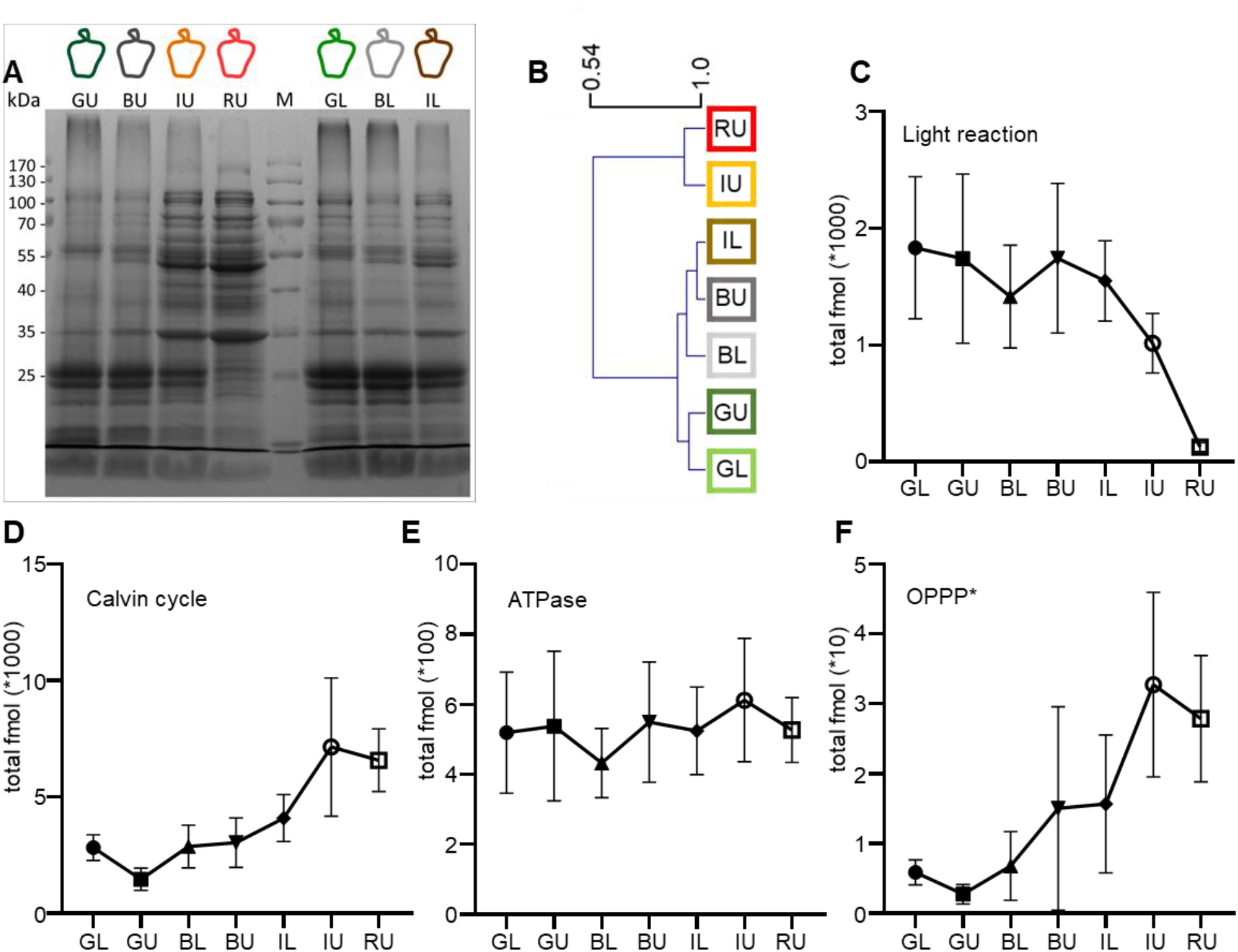
Protein profiles and protein abundances in plastids from different ripening stages. (A) SDS PAGE displaying protein profiles obtained from the different development stages during fruit ripening (See Fig. 1B) (B) Results of a hierarchical cluster of the different plastid types (“sample” clustering) with Pearson correlation as distance metric. (C-F) Total abundance of proteins involved in the different pathways/functional categories of primary energy metabolism in the different plastid preparation, i.e. light reactions of photosynthesis encompassing photosystem I and II and LHC proteins (C), Calvin cycle enzymes (D), ATPase subunits (E) and the oxidative branch of the pentose phosphate pathway (F) (* glucose-6-phosphate 1-dehydrogenase (A0A1U8HAS8, A0A1U8GPM1) and 6-phosphogluconate dehydrogenase (A0A2G2Y8Q9, A0A2G2Y8R9, A0A1U8F1N4)).

We used the quantitative proteome data to assess the similarity between the different plastid types by hierarchical clustering using a Pearson correlation distance metric (Fig. 3B). The plastid types cluster into two main cluster, one comprising the stages GL, GU, BL, BU and IL and the second comprising the IU and the RU stages. Within each of the two cluster, the plastid types are very similar with Pearson correlation coefficients > 0.8 (Fig. 3B). Surprisingly, the two intermediate stages IL and IU are clearly distinct despite having the same age (47 days, see Fig. 1B for reference), suggesting that the intermediate stage of fruit ripening comprises two different plastid types within one fruit. This is similar to the situation found in tomato, where the intermediate, yellow ripening stage contained chloroplasts and chromoplasts at the same time (Camara et al., 1995). In contrast to tomato, the two plastid types are not interspersed in pepper but rather confined to different sections in the fruit.

The most significant adaptation of the proteome to a heterotrophic chromoplast metabolism is the disassembly of the photosynthetic machinery as shown in Fig. 2 and in Fig. 3C. Notably, the protein constituents of the light reactions of photosynthesis are maintained up to the IL stage before they decline rapidly in the IU and RU stages (Fig. 3C). In contrast, constituents of the Calvin cycle and ATPase are maintained throughout the entire differentiation process. In case of ATPase this suggests, that fully mature chromoplasts synthesize ATP at high rate (see also discussion below, Fig. 3D and E). Similar to other heterotrophic plastid types, the oxidative branch of the pentose phosphate pathway (OPPP) is significantly elevated in chromoplasts (Fig. 3F), supporting the previous notion that a substantial amount of reducing power in heterotrophic plastids is generated by NADP reduction through OPPP activity (Neuhaus and Emes, 2000, von Zychlinski et al., 2005).

High OPPP activity in chromoplasts can be also inferred by the central role of ferredoxin (Fd) in the bell pepper chromoplast redox metabolism (Fig. 4). The abundance of ferredoxin, Fd-dependent NADPH reductase (FNR) and Fd-dependent glutamate synthase increases in the course of chromoplast differentiation (Fig. 4), which is counterintuitive because ferredoxin reduction relies on photosynthetic electrons that are not available at high supply in chromoplasts. It was demonstrated that root-type FNRs are able to reduce ferredoxin with electrons from NADPH (Green et al., 1991) suggesting that ferredoxin reduction in chromoplasts occurs with electrons from NADPH through activity of a root type FNR. Indeed, one out of two identified chromoplast FNRs (A0A1U8FJG3) has 79% identity to Arabidopsis root-type FNR1 (AT4G05390). Thus, NADPH generated through OPPP activity reduces ferredoxin, thereby feeding electrons into amino acid biosynthesis through Fd-dependent glutamate synthase (Fig. 4). Amino acid biosynthesis is a dominant metabolic activity in chromoplasts (Supplemental Table S3) as well as in other heterotrophic plastid types (Neuhaus and Emes, 2000, Baginsky et al., 2004, Siddique et al., 2006, von Zychlinski et al., 2005). Amino acid profiling at the onset of ripening reported a clear increase of 18 analyzed amino acids (Osorio et al., 2012) thus documenting the requirement for high amino acids biosynthetic activities.

**Figure 4:**
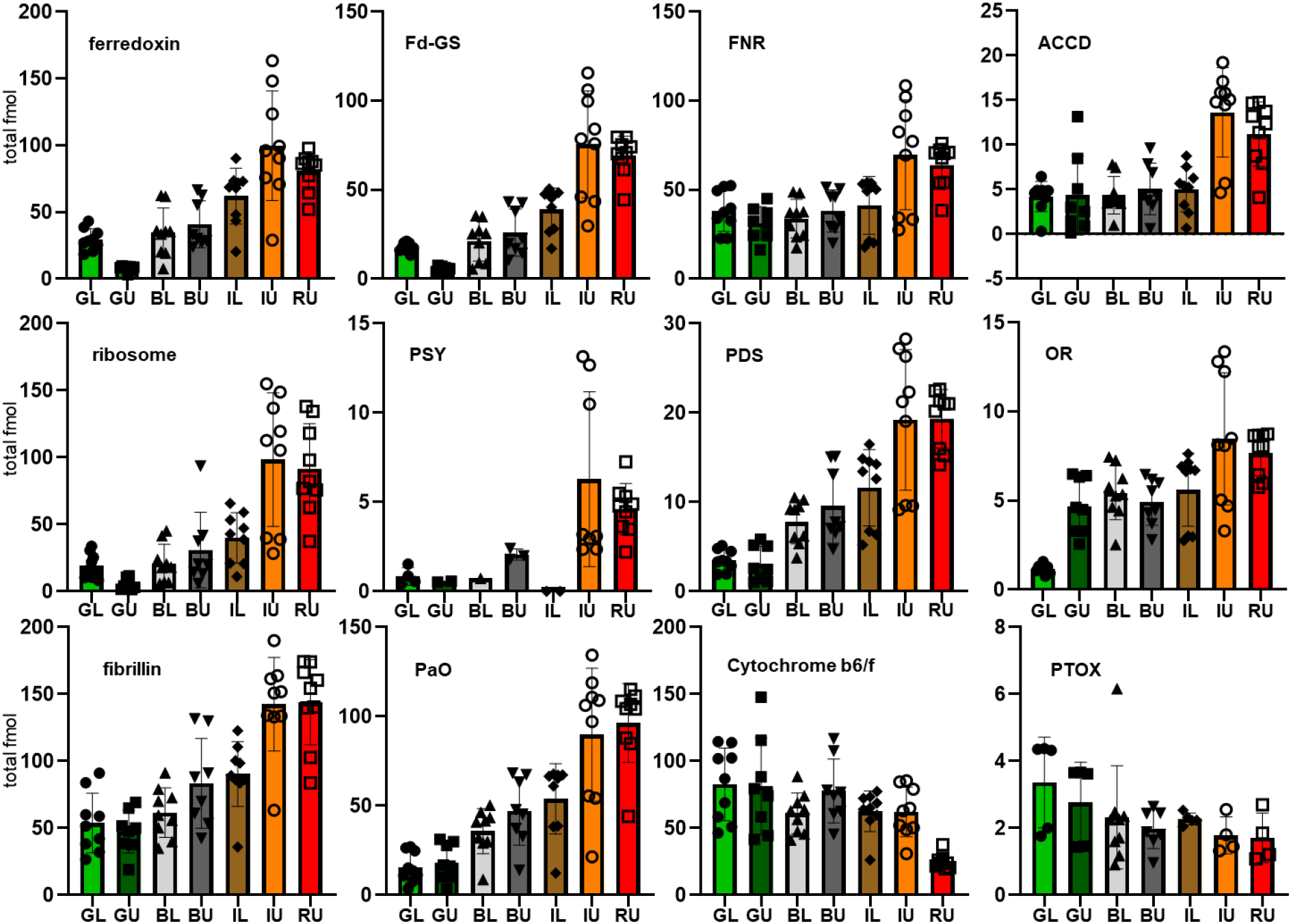
Abundance of selected proteins as determined from the in solution digests with the different plastid preparations. Provided are fmol abundances from three biological replicates (three technical replicates each) along with the standard deviation (SD) of the repeat measurements and the individual data points. The plastid preparations are represented by their abbreviation as in figure 1 (Fig. 1) and additionally color-coded. Following proteins/enzymes are presented: ferredoxin (A0A1U8GXA1, A0A1U8H397, A0A2G2YH86, A0A1U8DVL9, B1PDK3, A0A2G2XUS9), Fd-dependent glutamate synthase (Fd-GS - A0A2G3ANJ0, A0A2G2ZX64), Fd-dependent NADP reductase (FNR-A0A1U8FJG3, A0A1U8FMQ7), acetyl CoA-carboxylase (ACCD-A0A2G2ZQ00, A0A2G2Z7T4), plastid 70S ribosome (ribosome – see Supplemental Table S3), phytoene synthase (PSY-A0A1U8FLW3, A0A1U8GGI5), phytoene desaturase (PDS-A0A1U8FU73), ORANGE (OR-A0A1U8FXL9, A0A2G2ZY11), fibrillin (Q42493), pheophorbide a oxygenase (PaO-A0A2G2YHJ4), cytochrome b6/f complex (see Supplemental Table S3) and plastid terminal oxidase (PTOX-A0A2G2Y4F7).

High carotenoid biosynthesis is a hallmark of bell pepper fruit ripening. Carotenoids are sequestered to membranes; thus high fatty acid biosynthetic activity is necessary to sustain membrane biogenesis to accommodate storage carotenoids. The plastid encoded acetyl Co-A carboxylase subunit AccD is significantly upregulated during chromoplast differentiation (Fig. 4). AccD is essential for chloroplast fatty acid metabolism since it catalyzes the first committed step of fatty acid biosynthesis i.e. carboxylation of acetyl Co-A to malonyl Co-A. This plastid encoded protein is considered to have the highest translation rate in chromoplasts of tomato (Kahlau and Bock, 2008). Our proteomics data show that AccD is only moderately more abundant in chromoplasts compared to chloroplasts, while other plastid-encoded protein, e.g. components of the plastid ribosome remain abundant in chromoplasts as well (Fig. 4). The upregulation of fatty acid synthesis is paralleled by an increase of proteins involved in carotenoid biosynthesis as exemplified for phytoene synthase (PSY) and phytoene desaturase (PDS) and the DnaJ-like chaperone ORANGE (OR) (Fig. 4).

Plastoglobules are important for carotenoid sequestration (reviewed in van Wijk and Kessler, 2017). We find a 32 kDa fibrillin significantly upregulated during chromoplast differentiation, consistent with its function in plastoglobule structural organization and chromoplast pigment accumulation (Fig. 4). Recently, identification of ζ-carotene desaturase, lycopene β-cyclase and β-carotene β-hydroxylases, operating in plastoglobuli of chromoplasts suggested that plastoglobules function as both, lipid biosynthesis and storage compartments (van Wijk and Kessler, 2017). The investment in structures for carotenoid sequestration goes together with increasing abundance of chlorophyll degrading enzymes, as exemplified for pheophorbide a oxygenase (PaO) (Kuai et al., 2018) (Fig. 4).

### Chromorespiration

As in previous proteome analyses with tomato, our data show that the chloroplast ATPase is maintained at high level, similar to the cytochrome b_6_f complex (Fig. 3E and Fig. 4) (Barsan et al., 2012). Several proteins involved in the thylakoid electron transport chain and ATP synthesis, e.g. cytochrome b_6_f complex, PTOX, and NAD(P)H dehydrogenases were previously considered to contribute to chromoplast ATP synthesis (Pateraki et al., 2013, Renato et al., 2014). It is conceivable that electrons are transferred into the chromoplast membrane system at high rate, because phytoene desaturase catalyzes two consecutive dehydrogenation reactions with phytoene, transferring the electrons to PQ (Nievelstein et al., 1995; Norris et al., 1995). Thus, with the dramatic increase in phytoene synthase and phytoene desaturase observed here (Fig. 4), a massive transfer of electrons into the PQ pool is likely to occur during carotenoid biosynthesis. A recent model suggested that these electrons are scavenged by PTOX, that transfers excess electrons to O2 giving rise to H2O (Josse et al., 2000). This model is mostly based on the observation, that PTOX is present in fully differentiated chromoplasts and its abundance increases during tomato chromoplast development. It was shown, that PTOX activity is responsible of one quarter of total fruit oxygen consumption in fully ripe tomato fruits (Shahbazi et al., 2007, Renato et al., 2014, Nawrocki et al., 2015).

A similar conclusion has been reached for bell pepper since the expression of PDS, ZDS and PTOX is co-regulated at the transcriptional level (Renato et al., 2015). Our data show, however, that PDS and PSY abundance on the one hand, and PTOX abundance on the other are uncoupled at the protein level. While PDS and PSY abundances increase significantly as does the abundance of other carotenoid biosynthetic enzymes, PTOX abundance is rather constant during chromoplast differentiation (Fig. 4). Thus it is possible that the electrons from phytoene could pass through the cytochrome b_6_f complex adding a supplementary proton pumping site and being used by a currently unidentified oxidase. This hypothesis is supported by the effect of a cytochrome b_6_f inhibitor on chromoplast ATP synthesis (Renato et al., 2014). To identify proteins that are closely co-regulated with phytoene desaturase, we performed hierarchical clustering using a Pearson correlation distance metric. This cluster results in one cluster branch that contains a set of enzymes highly up-regulated in the course of chromoplast differentiation (Fig. 5). It comprises carotenoid biosynthetic enzymes, enzymes involved in fatty acid metabolism, the chaperone ClpB3, the plastid division protein FtsZ, the chlorophyll degrading enzyme pheophorbide a oxygenase and fibrillin (Fig. 5). Thus, this cluster branch concisely summarizes key features of chromoplast differentiation (see discussion above), but did not identify a previously unknown oxidase that may serve as the electron acceptor site required for phytoene desaturation.

**Figure 5:**
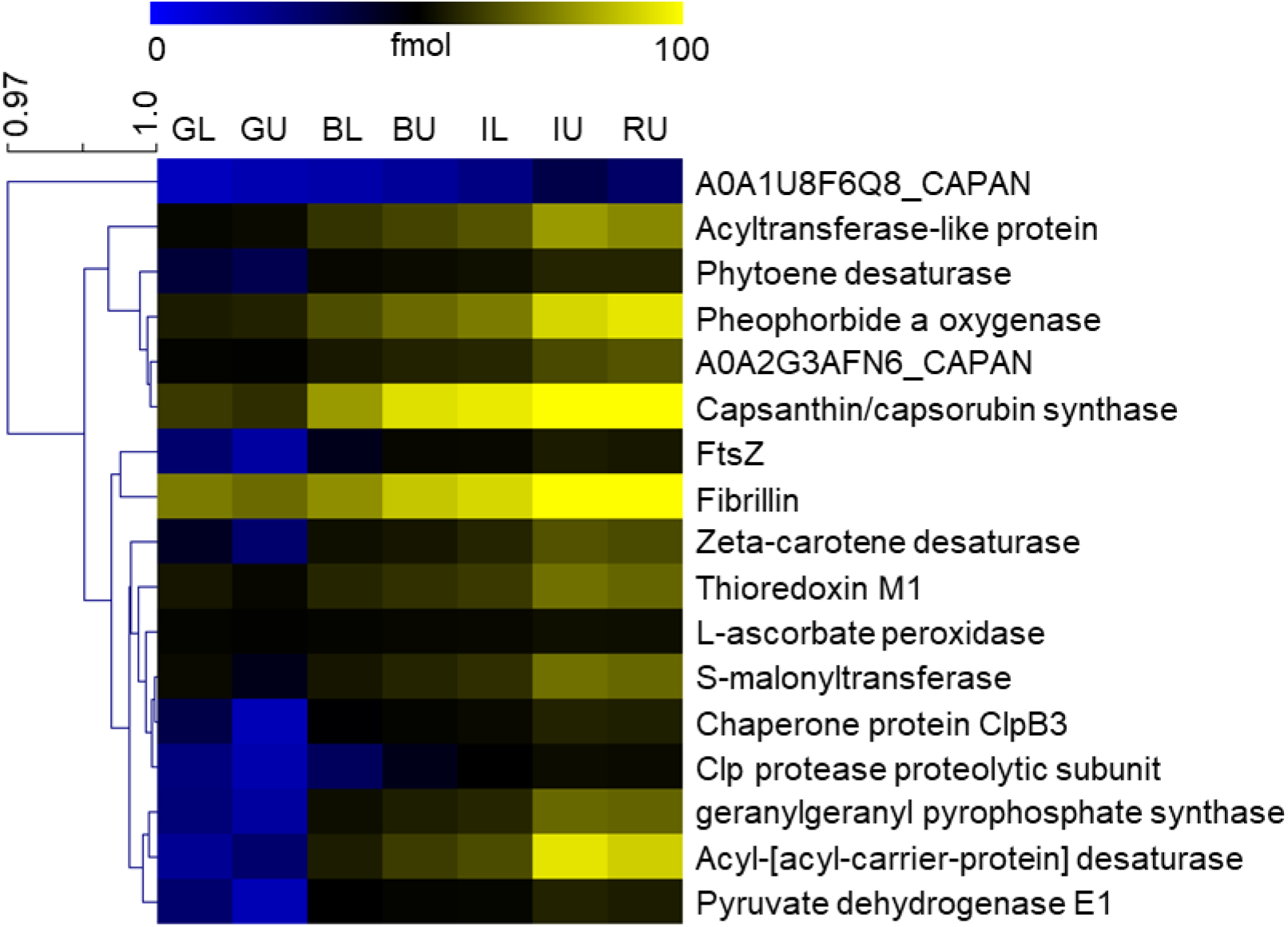
Hierarchical cluster around phytoene desaturase. We performed hierarchical clustering with all identified chromoplast proteins to identify co-regulated functional modules using a Pearson correlation distance metric. The presented branch of the cluster shows those proteins that are tightly co-regulated with phytoene desaturase in the course of chromoplast differentiation.

### Organization of the protein import machinery during chloroplast to chromoplast differentiation

The plastid protein import machinery is essential for chloroplast biogenesis and recent data advocated its reorganization during plastid differentiation to accommodate the changing requirements on import specificity originating from new import cargo. Intriguingly, the ubiquitin proteasome system (UPS) is involved in this process by removing *translocon at the outer chloroplast membrane* (TOC) components that are labeled for degradation by the E3 ubiquitin ligase Sp1, thus enabling their replacement by other TOC subunits with different precursor specificities (Sadali et al., 2019). While comprehensive analyses have questioned a clear substrate specificity of individual TOC subunits (Bischof et al., 2011, Dutta et al., 2014, Grimmer et al., 2020), the importance for Toc159 during de-etiolation and seed germination was clearly demonstrated (Ling et al., 2012, Shanmugabalaji et al., 2018).

The bell pepper Toc159 homolog was identified at low concentrations in only one to two replicates in the GL and GU samples, thus hampering its quantification. In contrast, the Toc75 and Toc33/34 homologs were identified at all stages during chromoplast differentiation (Fig. 6A). Their abundance is slightly higher in the IU and RU samples compared to the other plastid preparations (Fig. 6) supporting conservation of the TOC machinery in chromoplasts. A better coverage of TOC subunits was achieved in the CN-PAGE analysis (Supplemental Table S4). Here, Toc159, Toc75 and Toc33/34 were identified reproducibly in the high molecular mass region of the gel (Gel slices CN1 - CN3, Fig. 6B, Schäfer et al., 2019). In these complexes, the ratio between Toc159 and Toc75 is not altered between chloroplasts and chromoplasts thus there is no indication for a reorganization of the TOC apparatus. A comparison of transit peptides revealed no difference in the major amino acid composition of the 100 most abundant proteins in GL and the RU plastids (Supplemental Fig. S3), thus there may not be major shifts in specificity required for the import of chromoplast proteins. Nonetheless, we cannot exclude that TOC complexes with the bell pepper homologs of Toc132 or Toc120 may be more abundant in chromoplasts compared to chloroplasts, since we did not detect any large GTPases other than Toc159 (Agne et al. 2009). Thus, while it is clear that Toc159 is incorporated in high molecular mass TOC complexes in chromoplasts (Fig. 6B), it remains unclear whether chromoplast protein import is additionally sustained by alternative TOC complexes.

**Figure 6:**
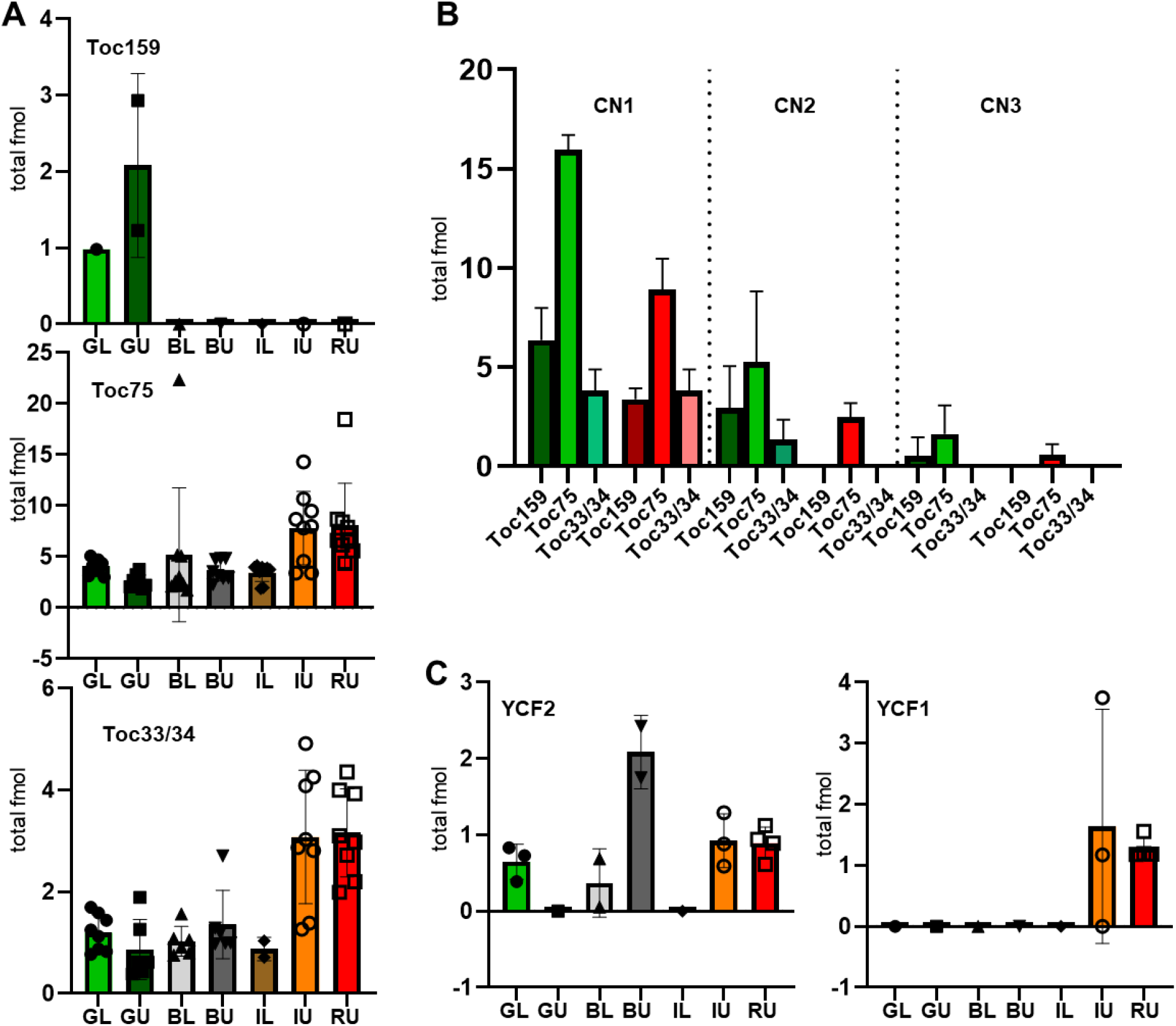
Abundance of subunits of the translocases at the outer (TOC) and inner (TIC) envelope membranes of chloroplasts. The abundance of the bell pepper homologs for Arabidopsis Toc159 (A0A2G2ZW43) (note that the abundance value was determined from only one (GL) or two (GU) data points and as such is unreliable), Toc75 (A0A2G2ZEA8) and Toc33/34 (A0A1U8G0U4) as identified from *in solution* digests (A) and from the CN-PAGE analyses (B). CN1-3 – CN PAGE gel slice 1-3 (see also fig. 2) (C) The abundance of the inner envelope translocase components YCF1 (A0A2G2YAF6) and YCF2 (A0A2G2YSI8, A0A2G2Z199, A0A2G2ZTJ0, A0A2G3AIN8) as determined from in solution digests. In all instances error-bars display standard deviation (SD).

It was suggested that the plastid gene expression machinery is maintained mainly in order to produce AccD (see above). However, recent results suggested that YCF1 and YCF2 are essential for plastid functioning because both are components of the protein import machinery, YCF1 as component of a 1-MDa translocase at the inner envelope membrane and YCF2 as component of a 2-MDa ATP hydrolysis activity associated with the TIC translocon (Kikuchi et al., 2013, Kikuchi et al., 2018). The protein import machinery is maintained at high levels to support the re-organization of the proteome during chromoplast development (Fig. 6 A and B). This encompasses e.g. the massive import of carotenoid biosynthetic enzymes that are, without exception, nuclear-encoded. With the exception of low amounts of plastid encoded YCF1 and YCF2, no other inner envelope translocase (TIC) subunits were identified (Fig. 6 C).

## Concluding remarks

Our quantitative proteome analysis reported here provides a fine-grained documentation of the chromoplast differentiation process in bell pepper fruits. Earlier quantitative proteome analyses of the chromoplast differentiation process were conducted with the climacteric fruit tomato. Our analysis with bell pepper therefore represents the first dataset for chromoplast differentiation in a non-climacteric fruit. Our data confirm some observations made in tomato such as changes in the amount of enzymes involved in carotenoid biosynthesis, tetrapyrrole biosynthesis and chlorophyll degradation. While photosynthetic proteins progressively decline in abundance, typical enzymes of the heterotrophic plastid metabolism such as fatty acid and amino acid biosynthesis and those of the oxidative branch of the pentose phosphate pathway (OPPP) increase in abundance. In contrast to the tomato data, we do not find an increase in plastid terminal oxidase levels (PTOX) and only a moderate increase in AccD abundance (Fig. 4). We also uncovered unexpected features of the redox metabolism in bell pepper chromoplasts, that appears to operate to a large extent via ferredoxin. With this manuscript we make all mass spectrometry data files available for download via PRIDE (https://www.ebi.ac.uk/pride/archive/), facilitating their re-analysis with different tools for database matching. In summary, our data provide insights into the chromoplast differentiation process at unprecedented depth and spark new hypotheses that await further testing.

## Acknowledgments

S.B. gratefully acknowledges DFG support for the acquisition of a Synapt G2-S mass spectrometer (INST 271/283-1 FUGG) and for support through DFG grant BA 1902/3-2. We furthermore sincerely thank Christian E. H. Schmelzer (Fraunhofer Institute for Microstructure of Materials and Systems (IMWS) Halle (Saale)) for the excellent cooperation in the analysis of pigments with mass spectrometry. The authors furthermore thank Arne Preuß for help with bell pepper growth.

## Materials and Methods

### Material

LC-MS grade solvents, including water with 0.01% (w/v) formic acid, water with 0.1% (w/v) trifluoroacetic acid and acetonitrile with 0.1% (w/v) formic acid were obtained from Carl Roth (Karlsruhe, Germany). Porcine sequencing grade modified trypsin was obtained from Promega (Mannheim, Germany).

### Plant Material

*Capsicum annuum* fruits were used for the preparation of chloroplast, chromoplast and pigment extracts. The plants (cultivar Selma Bell) were grown on soil in a greenhouse (20.8-15.7 °C, 16 h light, relative humidity: day 65%, night 45%). Fruits of different degrees of maturity (Figure 1A) were harvested immediately after the end of the dark period. 180-200 g of chopped plant material were homogenized in 360-400 mL homogenization buffer (330 mM sorbitol, 2 mM EDTA, 50 mM HEPES (pH 7.5), 1 mM MnCl_2_, 1 mM MgCl_2_, 2 mM DTT, 0.1% (w/v) BSA, 0.1% (w/v) Na ascorbat) and filtered through Miracloth (22-25 μm). The pellet (2000 × g, 4 °C, 10 min) was resuspended in 5 mL homogenization buffer and used for Percoll density gradient centrifugation (2.5 mL each, 40 min, 2600 × g, 4 °C, acceleration set 3, brakes set off). Upper (40/20%) and lower (60/40%) layers containing plastids were transferred into 25 mL HEPES (50 mM, pH 7.5) / sorbitol (330 mM) buffer and centrifuged (5 min, 2000 × g, 4 °C). The plastid pellet was washed once by adding 10 mL HEPES/sorbitol buffer. Plastid pellets were resuspended in HEPES/sorbitol buffer. Aliquots of 250/500 μg protein (Bradford) were frozen in liquid N2 and stored at −80 °C.

### Pigment analysis

Pigments of all different maturation stages (green, black, intermediate, red) were extracted using 500 μL MeOH:chloroform (2.5:2, v/v, 0.1% butylated hydroxytoluene) per 250 μg plastid pellet. Upon incubation (10 min, on ice in the dark) 250 μL Tris-HCl (pH 7.5, 1 M NaCl) were added. After 10 min on ice in the dark, samples were centrifuged (10 min, 1350 × g, 4 °C). Aqueous phase was re-extracted twice using 500 μL 0.1% BHT in chloroform and merged organic phases were dried in vacuum and stored until further analysis. UV/Vis absorption spectra were recorded between 220-800 nm and the samples were stored at −20 °C. In order to quantify changes during maturation, all samples were analyzed on a LaChrom HPLC (Merck-Hitachi: L-7100 pump, D-7000 interface, L-7455 DAD detector) equipped with a ProntoSIL C18 AQ (120 Å, 5 μm). 20 μL of a re-solubilized (in 150 μL acetone,) pigment samples were injected and separated in a linear gradient of 30-90% B (A: methanol:H2O (75:25, v/v), B: ethyl acetate) within 50 min. The eluent was detected for 70 min in the range of 230-750 nm. Pigments were identified by atmospheric pressure chemical ionization mass spectrometry on an LCQ Classic Finnigan MAT (fullscan, positive mode, 50-2000 m/z, 400 °C APCI, 7μA corona voltage, 250 °C capillary temperature) coupled to an Agilent 1220 Infinity LC, equipped with the above-mentioned RP18 column in a linear gradient of 30-80% B in 30 min and 80-90% B in additional 20 min (A: methanol:H2O (75:25, v/v), B: 2-propanol).

### Colorless Native PAGE

Aliquots of 250 μg protein were resuspended in 125 μL CN buffer (50 mM Bis-Tris (pH 7), 0.5 M ε-aminocaproic acid, 10% glycerin, 5 mM DTT, 2 mM CaCl2, 20 μL/mL protease inhibitor mix (Serva), 1,5% n-Dodecyl β-D-maltoside) to obtain a final concentration of 2 μg/μL protein or 0.5 μg/μL chlorophyll, respectively. Upon incubation (30 min, 4 °C, 15 rpm) and centrifugation (30 min, 100000 × g, 4 °C) 31.2 μL 5× CN buffer (50 mM Bis-Tris (pH 7), 0.5 M ε-aminocaproic acid, 50% glycerin, 0.04‰ Ponceau S,) were added to the supernatants. The dissolved samples were centrifuged (5 min, 20000 × g, 4 °C) and applied to gradient gels (5 - 13.5%). CN polyacrylamide gels were stained with Coomassie Brilliant Blue.

### Protein digest

Every CN polyacrylamide gel lane was cut into 16 equally-sized slices that were subsequently digested with trypsin as previously described (Rodiger et al., 2014). For the in solution digests, aliquots of 250 μg protein were resuspended in 100 μL 25 mM ammonium bicarbonate (final protein concentration 5 μg/μL). 20 μL (100 μg protein) were solubilized (10 min, 80 °C) in 201 μL 25 mM ammonium bicarbonate containing 0.05% (w/v) RapiGest SF (Waters), and reduced by adding 6.2 μL 10 mM dithiothreitol (10 min, 60 °C). Cystein residues were alkylated using 6.2 μL 30 mM iodacetamide for 30 min in the dark. Subsequently, 2 μL trypsin (final concentration: 1:100 (w/w) corresponding to 1 μg protease per vial) was added. Samples with a final volume of 250 μL were digested at 37 °C overnight. To stop the digestion, 2 μL 37% HCl was added to reach a pH value < 2. To avoid loading insoluble material onto the column, the peptide solutions were thoroughly centrifuged prior to sample loading (4 °C, 21500 × g, 30 min).

### Nano-LC Separation, HD-MS^E^ Data Acquisition and Protein Identification/Quantification

Nano-LC-HD-MS^E^ data were acquired for three biological replicates and for every biological replicate, three technical replicates were performed. Data acquisition was performed as described previously (Helm et al., 2014) using a nanoACQUITY UPLC (trap column: 200 mm × 180 μm fused silica, 5 μm Symmetry C18; separation column: 250 mm × 75 μm fused silica, 1.8 μm HSS T3 C18) / Synapt G2-S mass spectrometer (Waters). Data analysis was carried out by ProteinLynx Global Server (PLGS 3.0.1, Apex3D algorithm v. 2.128.5.0, 64 bit, Waters) with auto-determination of chromatographic peak width and MS TOF resolution. Lock mass window was set to 0.25 Da. Low/high energy threshold was set to 180/15 counts, respectively. Elution start time was 5 min, intensity threshold was set to 750 counts. Databank search query (PLGS workflow) was carried out as follows: Peptide and fragment tolerances was set to automatic, at least 2 fragment ion matches per peptide and 5 fragment ion matches from at least 2 peptides were required for protein identification. Maximum protein mass was set to 250 kDa. Primary digest reagent was trypsin (1 missed cleavage). According to the digestion protocol fixed (carbamidomethyl on Cys) as well as variable (oxidation on Met) modifications were set. 10 fmol/injection rabbit glycogen phosphorylase B (P00489) was defined as internal quantification standard. The database for bell pepper (*Capsicum annuum*) was downloaded from https://www.uniprot.org/proteomes/UP000222542, containing 35,548 protein entries (Qin et al., 2014). The false discovery rate (FDR) was 4% at protein level. The mass spectrometry proteomics data have been deposited to the ProteomeXchange Consortium (Vizcaíno et al., 2016) via the PRIDE partner repository (https://www.ebi.ac.uk/pride/archive/) with the dataset identifier PXD006409 and PXD006477.

### Hierarchical clustering

Clustering analyses were performed using the Multi Experiment viewer (http://mev.tm4.org/). For the hierarchical cluster, we loaded the average abundance values for every protein in the different plastid preparations into the data file, and performed clustering with a Pearson correlation metric using average linkage clustering.

## Supplemental Figures

**Supplemental Figure 1:**
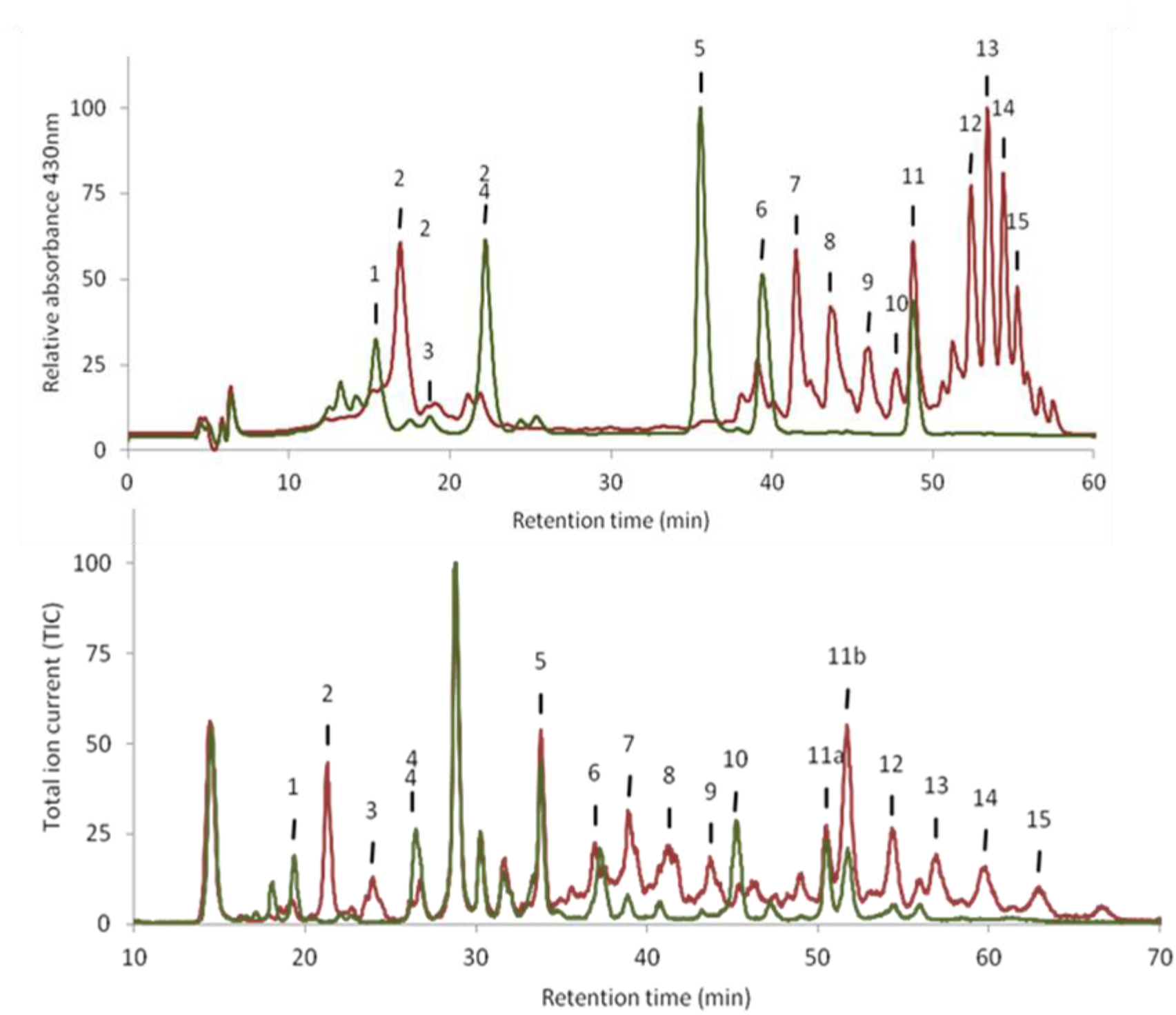
Chromatograms of pigment extracts of chloroplasts (green) and chromoplasts (red) measured at 430 nm. Fifteen (15) peaks were numbered in order of their occurrence. For peak identity see Supplemental Table S1.

**Supplemental Figure 2:**
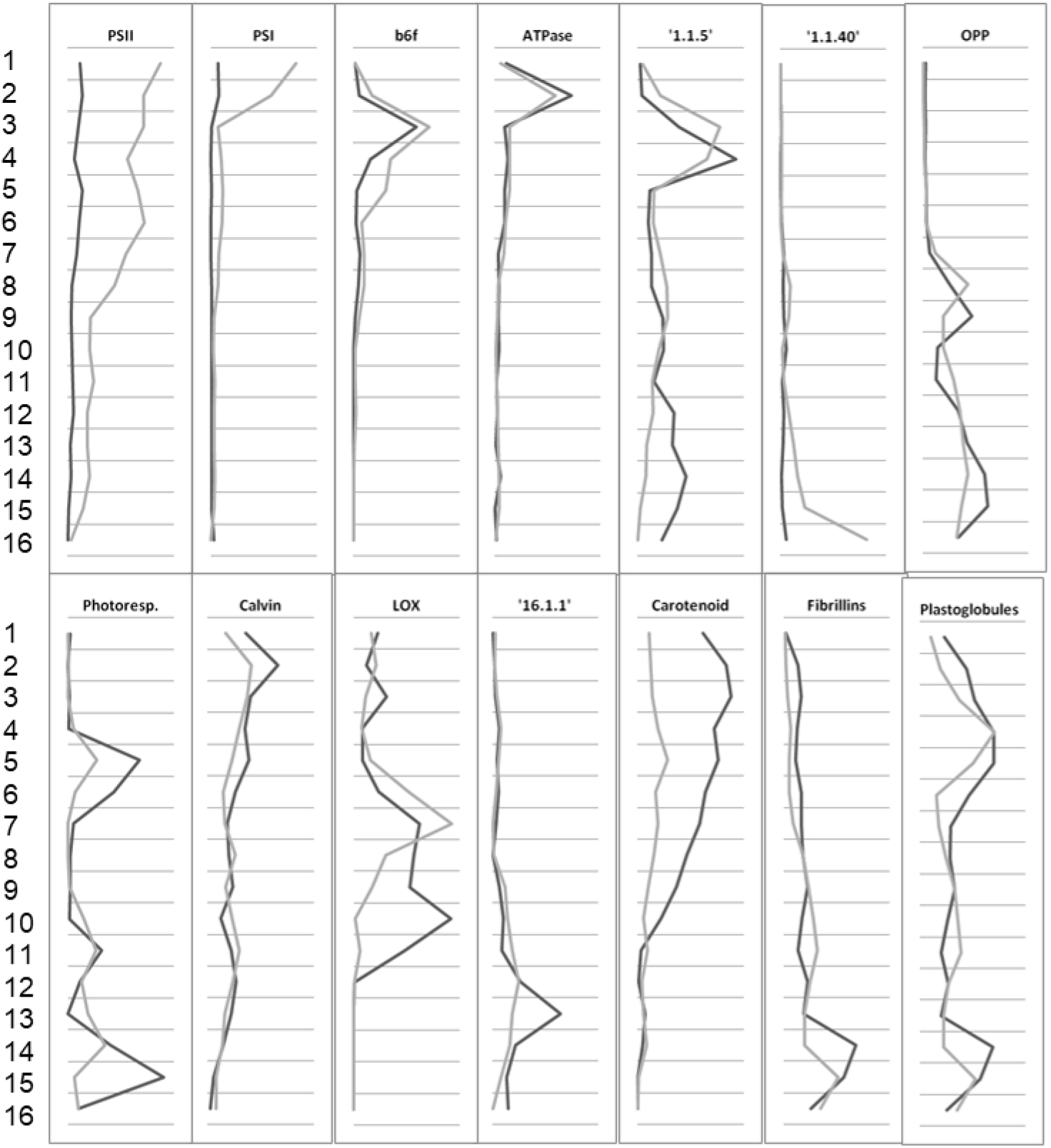
Profile of protein abundances along the CN-PAGE gel in every gel slice (compare to Fig. 2, light gray chloroplast (GL), dark gray chromoplasts (RU)). Provided are the abundances for the two photosystems PSI and PSII, the cytochrome b6/f complex, ATPase, MapMan bins 1.1.5 and 1.1.40, oxidative pentose phosphate pathway (OPPP), photorespiration, Calvin cycle, lipoxygenases (LOX), MapMan bin 16.1.1, carotenoid synthesis, fibrillins and plastoglobules. Data were matched against the Arabidopsis database and functional annotation was based on MapMan (Thimm et al., 2004).

**Supplemental Figure 3:**
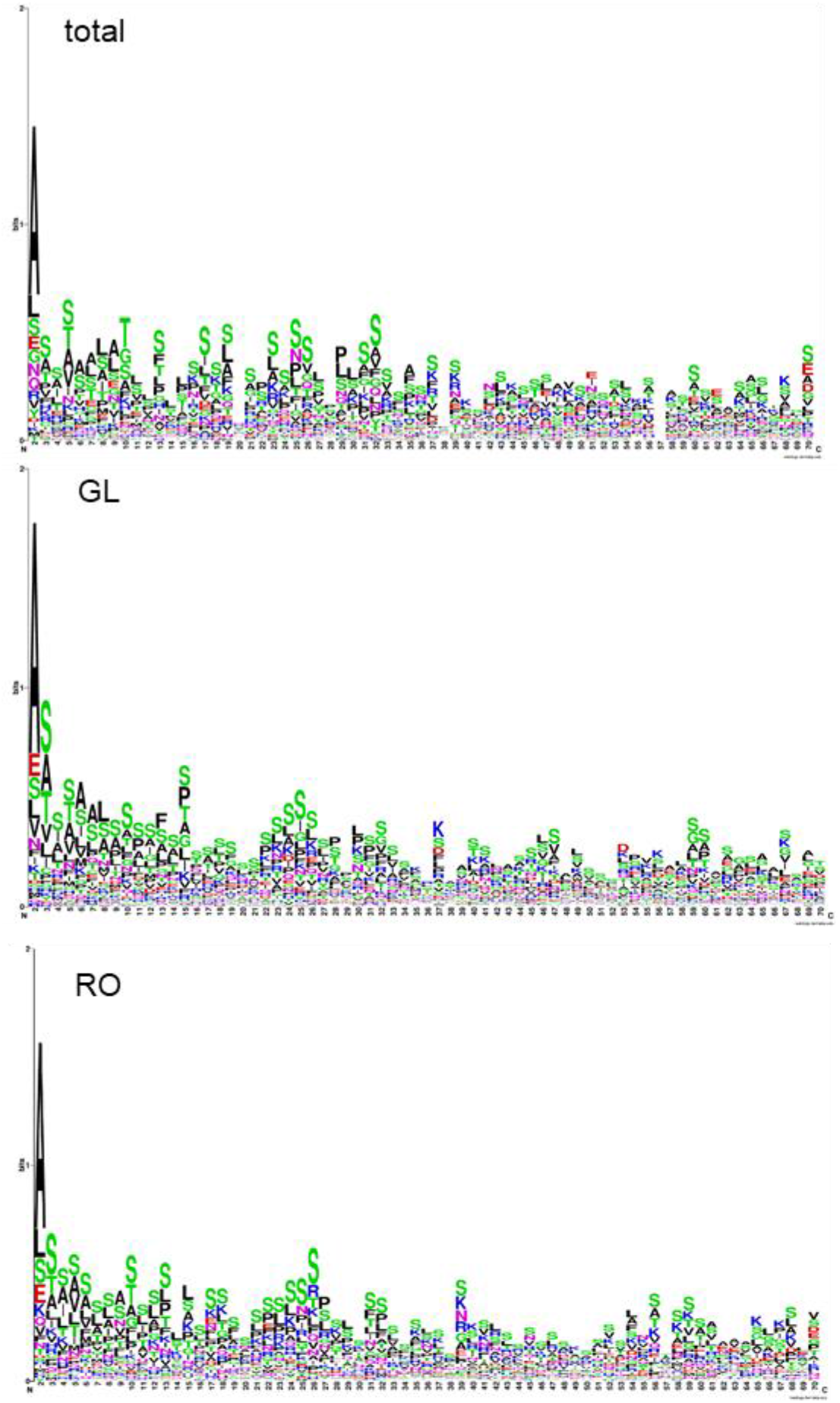
Alignment of putative transit peptides of chloroplast and chromoplast proteins. For the purpose of this alignment we extracted the 100 most abundant proteins from every plastid type isolated during this study and generated a composite list with non-redundant entries (total). These were aligned using WebLogo. To identify specific features of transit peptides in chromoplast proteins, we compared the alignment of the 100 most abundant chloroplast proteins (GL) with the alignment of the 100 most abundant chromoplast proteins. With this procedure, we intended to identify putative motifs in transit peptides that may require different import specificities among the TOC receptors, particularly with respect to a massive reorganization of the plastid proteome during the transition of different plastid types. No such motifs were detected.

